# The Effect of Anesthesia Gases on the Oxygen Reduction Reaction

**DOI:** 10.1101/2022.11.29.518334

**Authors:** Anu Gupta, Yutao Sang, Claudio Fontanesi, Luca Turin, Ron Naaman

## Abstract

The oxygen reduction reaction (ORR) is of high importance, among others because of its role in cellular respiration and in the operation of fuel cells. Recently, a possible relation between respiration and general anesthesia has been found. This work aims to explore whether anesthesia related gases affect the ORR. In ORR, oxygen which is in its triplet ground state is reduced to form products that are all in the singlet state. While this process is “in principle” forbidden because of spin conservation, it is known that if the electrons transferred in the ORR are spin polarized, the reaction occurs efficiently. Here we show, in electrochemical experiments, that the efficiency of the oxygen reduction is reduced by the presence of general anesthetics in solution. We suggest that a spin-orbit coupling to the anesthetics depolarizes the spins. This causes both a reduction in reaction efficiency and a change in the reaction products. The findings may point to a possible relation between ORR efficiency and anesthetic action.

**TOC:** 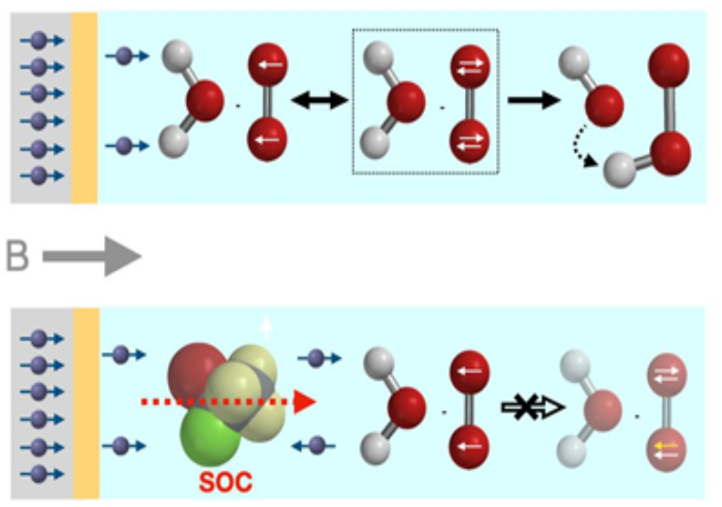

---

The oxygen reduction reaction (ORR) involves a reduction of oxygen by two pairs of electrons and it depends on the pH.^1,2^ The reaction is of high importance since it is responsible for the cellular respiration process and the functioning of fuel cells.^3^ In both of these applications, the oxygen, which is in a triplet ground state, reacts to form products that are in their singlet state. As is well established in chemical reactions, when a reagent is in a triplet state, most probably the product will also be a triplet. In the past, this “anomaly” in the efficiency of the respiration process was explained by large spin orbit coupling in the enzymes involved.^4^ This coupling can “mix” singlet and triplet states and thereby allows the process to occur at lower activation energies. Many of the enzymes have heavy atoms and metals that have large spin-orbit coupling; thus, they can mix singlet and triplet efficiently. However, some of the respiration-related enzymes do not contain heavy atoms and hence, it was not clear how they can facilitate the respiration process.^5,6^ In the case of fuel cells, expensive heavy metal catalysts must be used to provide the large spin orbit coupling.

In a recent study,^7^ it was shown that when the electrons that reduce the oxygen, have the same spin, the oxygen reduction reaction occurs more efficiently, compared to when the spins of the electrons are randomly oriented (see Figure 1). It is also known that when electrons pass through chiral molecules, their spins become aligned parallel to each other. This effect, known as chiral-induced spin selectivity (CISS), was found to occur when electrons are transmitted through various biomolecules and proteins.^8^ It was also shown that when the electrode is coated with chiral molecules, or is made chiral, the ORR reaction efficiency increases.

**Figure 1:**
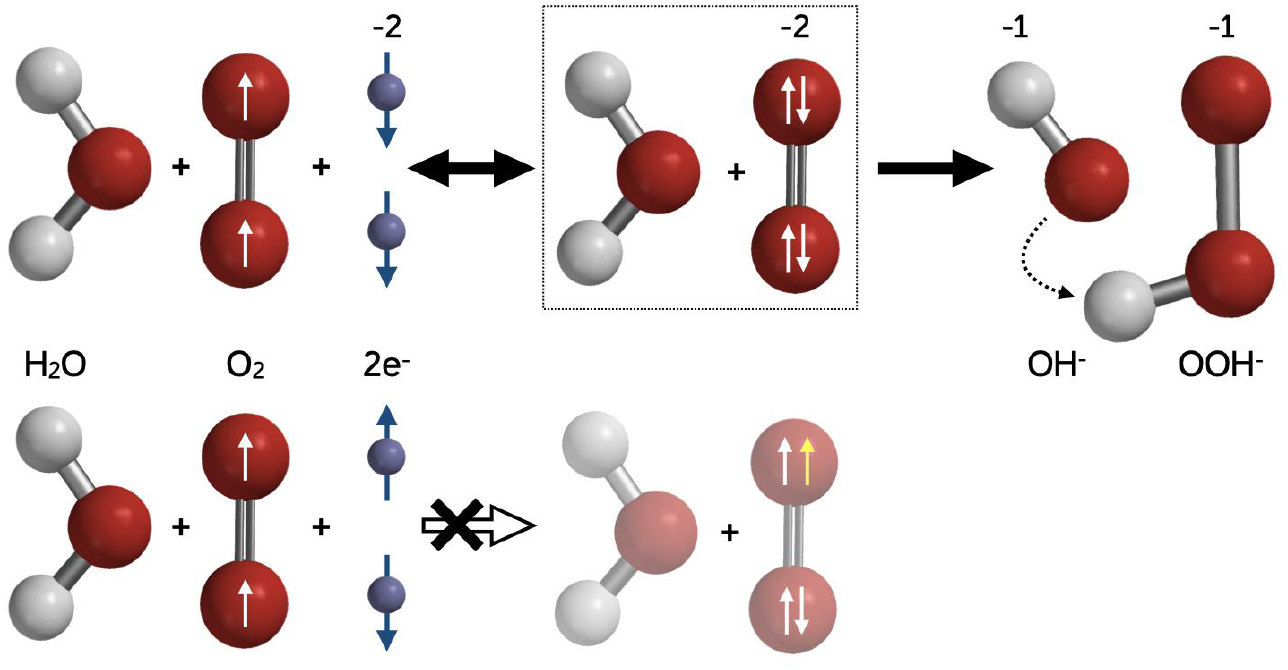
Effect of spin polarization on the oxygen reduction reaction (ORR) in alkaline aqueous media. **Top line**: Dioxygen in its triplet state reacts with spin-polarized electrons to yield hydroxide and hydroperoxide ions via a postulated dianion intermediate. **Bottom line**: The same reaction with unpolarized electron spins is forbidden.

Recently a compelling link between general anesthesia and respiration has been proposed, suggested by evidence of the involvement of mitochondrial proteins.^9,10,11^ Hence, the question we aimed to answer is how the anesthetic affect the ORR reaction, which is the last, crucial step in the respiration process. To answer this question we performed the ORR in an electrochemical cell (see Figure 2A) using two configurations. In the first, the reactions were investigated with a ferromagnetic working electrode made from silicon coated with a ferromagnetic Ni (150 nm)/Au (8 nm) layer. The reactions were performed with the electrode magnetized out of plane or with the electrode unmagnetized. As shown in the supporting material (Figure S5) when no external magnetic field is applied, the electrode is not magnetized; however, the magnetization is saturated at an applied field of about 0.42 Tesla. When the electrode is magnetized, the spins of the electrons ejected from it are mainly aligned parallel to each other and they occupy mainly one spin state.^12^ In the case of an unmagnetized electrode, the electrons ejected can be in either of the two spin states. We then added various gases to the reaction and monitored how they affect the reaction.

**Figure 2:**
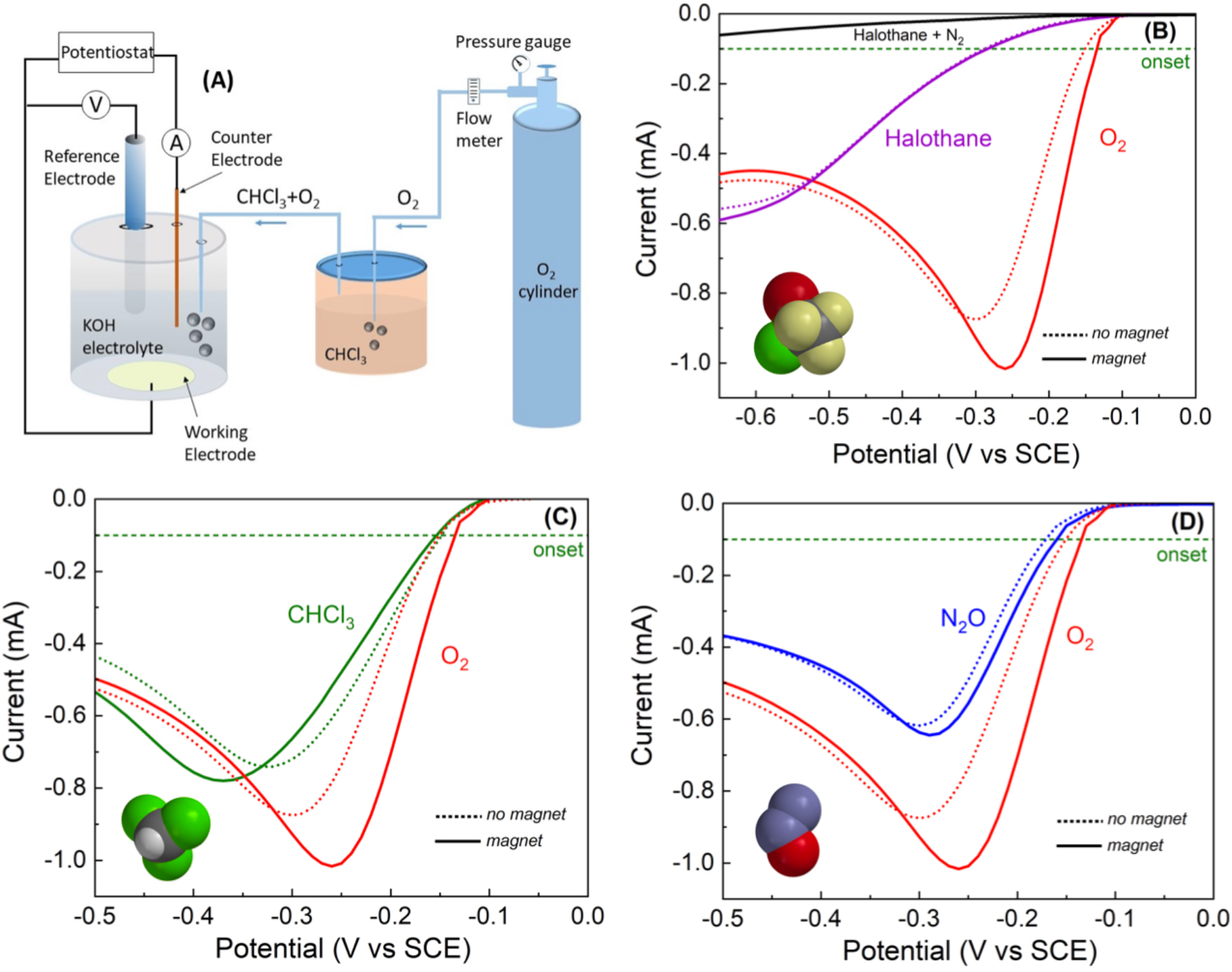
The electrochemical system and the results obtained for oxygen reduction. A: Schematics of the experimental set-up. B, C and D: The current versus the potential measured in the electrochemical system for magnetic (M-solid lines) and non-magnetic (NM-dotted lines) electrodes when anesthesia gases are added to the oxygen. B: halothane (2-Bromo-2-chloro-1,1,1-trifluoroethane) C: chloroform and D: nitrous oxide

In the second configuration, we used a working electrode made from “chiral gold”.^13^ The chiral gold is produced by electrodeposition of gold film from a solution containing a gold salt and an enantiopure chiral molecule. The electrons ejected from the chiral gold are spin polarized, as previously reported.^7^ The spin polarization results from the CISS effect. Hence, the results from the first configuration, the magnetic electrode, should be consistent with the results obtained with the chiral electrode. The use of the second setup is aimed at confirming that the effect is not a result of simply applying the magnetic field, but it is consistent with the spin being polarized due to the surface being magnetized or being chiral, which also occurs in bio-systems that are chiral.^14^ Also in this case gases were added to the reaction mixture.

Figures 2B, C and D present linear sweep voltammetry (LSV) curves, using the first experimental configuration (Ni/Au electrode), when the electrode is either magnetized (*magnet*) or not magnetized (*no magnet*). (See figure S5 for the magnetization curve of the electrode). The electrochemical measurements were performed in a solution saturated with oxygen only and with the same solution in the presence of various molecules. The LSV curves are recorded in the 0 to −0.5 V potential range with a potential scan rate of 50 mV/s. The current peak shown in the LSVs, in agreement with the experimental data present in the literature, at about −0.35 V.^2,15,16,17,18^ In the case of the solution free of any other gases, a difference is found when comparing the LSV curves recorded with a magnetized electrode (solid line) and the unmagnetized electrode (dashed line). A decrease of about 20% is found in the current at the peak of the LSV when the electrode is not magnetized versus a magnetized one (Table 1). Examination of the LSV curves when the anesthetic gases are added to the solution (Figures 2B, C, D) shows a clear decrease in the peak current and a shift to a higher threshold potential. In addition, the differences between the curves obtained with the magnetized (M-solid line) versus the non-magnetized electrode (NM-dashed line) are smaller, both regarding the onset ORR potential and regarding the value of the current at the peak. The results obtained with halothane are especially interesting, since they show a dramatic drop in the reduction peak and a very significant shift to a higher potential. The black line, at the top, is the voltammogram taken with halothane in the absence of oxygen, confirming that the shifted peak is indeed due to oxygen reduction.

**Table 1:** Summary of the results obtained from the electrochemical studies on various anesthetics added to the oxygen solution. The gases that are considered anesthesia efficient are in blue. The results with KBr are in yellow.

| Compounds<br>in solution | Current at the peak (mA) | | Threshold potential<br>(-V) | | Potential at the peak<br>(-V) | | Variation<br>(%)<br>$\frac{I_M - I_{NM}}{I_M} \times 100$ |
| --- | --- | --- | --- | --- | --- | --- | --- |
| | NM ($I_{NM}$) | M ($I_M$) | NM | M | NM | M | |
| O<sub>2</sub> | 0.78±0.09 | 0.99±0.05 | 0.16±0.01 | 0.14±0.01 | 0.29±0.01 | 0.28±0.01 | 21.21±0.14 |
| N<sub>2</sub> | 0.46±0.08 | 0.61±0.12 | 0.18±0.02 | 0.15±0.01 | 0.29±0.02 | 0.27±0.01 | 24.59±0.20 |
| Ar | 0.51±0.06 | 0.59±0.05 | 0.18±0.02 | 0.14±0.01 | 0.32±0.01 | 0.27±0.01 | 13.56±0.11 |
| CHCl<sub>3</sub> | 0.78±0.09 | 0.84±0.02 | 0.15±0.01 | 0.16±0.01 | 0.33±0.01 | 0.35±0.01 | 7.14±0.11 |
| N<sub>2</sub>O | 0.61±0.01 | 0.64±0.04 | 0.18±0.04 | 0.16±0.02 | 0.33±0.06 | 0.29±0.02 | 4.69±0.05 |
| CCl<sub>4</sub> | 0.71±0.07 | 0.70±0.09 | 0.18±0.03 | 0.18±0.03 | 0.37±0.01 | 0.35±0.01 | -1.41±0.16 |
| Halothane | 0.49±0.05 | 0.52±0.02 | 0.30±0.05 | 0.30±0.02 | 0.58±0.01 | 0.58±0.01 | 5.77±0.07 |
| KBr | 0.73±0.05 | 0.83±0.07 | 0.19±0.03 | 0.19±0.03 | 0.34±0.03 | 0.35±0.01 | 12.05±0.12 |

Interestingly, the bromide ion, which, owing to its ionic charge is nonvolatile, shows the same effect as the gases we used (see figure S1). The effect of gases that are not efficient anesthetic is much smaller as shown in Table 1 and in the Supplementary Material.

As a control experiment, the electrochemical process was carried out with a gold working electrode (not magnetic), with and without a magnetic field applied. In this case no effect of the magnetic field was observed (Figure S2). Note that in this case the electrons ejected from the working electrode have a randomly oriented spin and the magnetic field of 0.5 T is several orders of magnitude too small to align the electrons at room temperature. Hence, we can conclude that the magnetic field does not induce the effect by itself, but rather that it results from spin polarization within the ferromagnetic electrode that causes spin-polarized electron current.

Several parameters can be derived from the LSV (see Table 1). The first is the magnitude of the current at the peak. This value is sensitive to the concentration of oxygen. Although the concentration of oxygen may vary, to some extent, with dilution by the gases, either by mixing or bubbling, our focus is on the relative effect of a magnetized vs unmagnetized electrode on the oxygen current peak. Since the solution is the same independently of electrode magnetization, the difference observed is due to the effect of the anesthetic on the spin polarization of the electrons. The second parameter is the threshold potential (onset potential) at which the current starts to increase. To avoid confusion, we decided to take the value of the potential at which the current reaches 0.1 mA as the threshold potential. However, the most interesting parameters are the differences in the current and the threshold potentials when the electrode is magnetized, compared to the values for an unmagnetized electrode. From the LSV curves, one can conclude that in the case of pure oxygen, the curves with the electrode magnetized appear at a lower potential compared with the unmagnetized electrode.

To confirm the role of the electron spin, in the effects described above, we used a chiral gold electrode (see circular dichroism, CD, in Figure 3A and UV-Vis in Figure S4) in the electrochemical process and compared the results to those obtained with an achiral electrode (Figure 3B).

**Figure 3:**
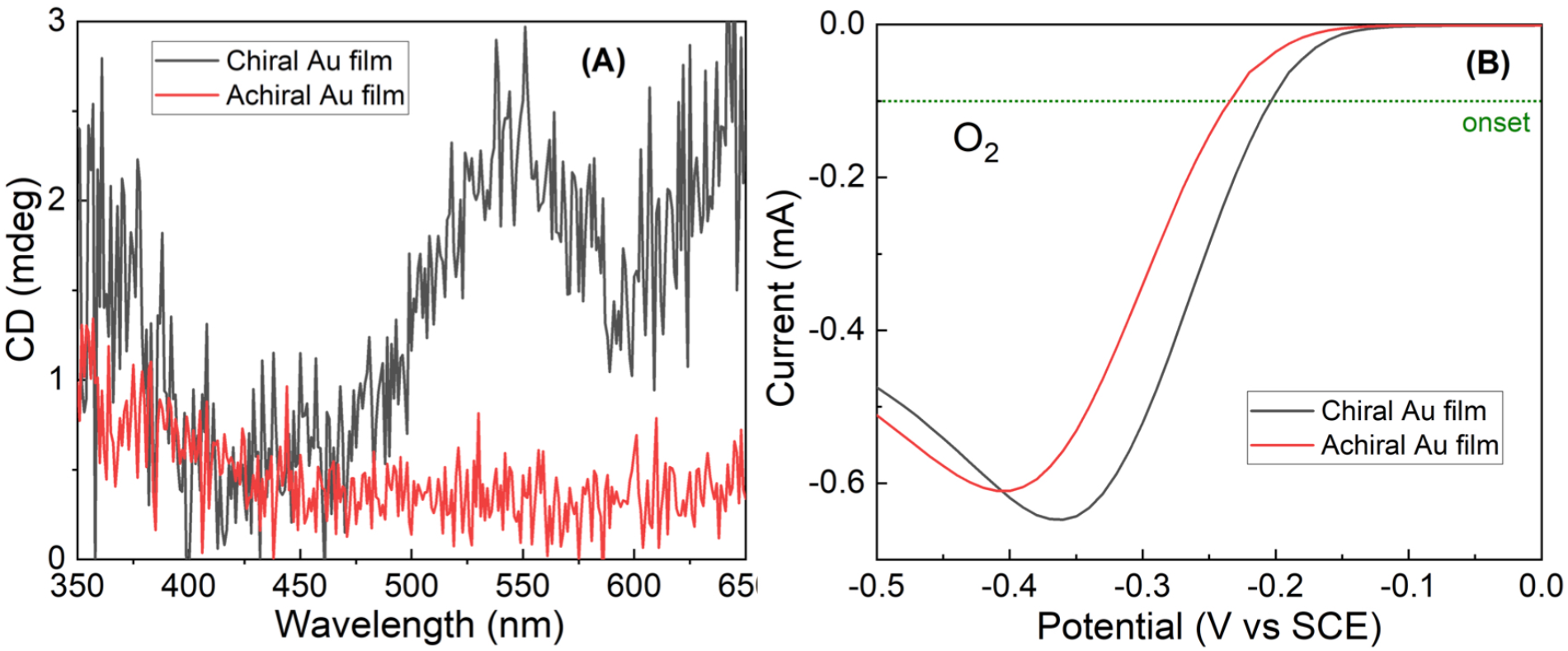
Oxygen reduction reaction with chiral and achiral gold electrodes. A) Circular dichroism (CD) spectra of the achiral and chiral electrodes. The CD signal appears only in the chiral electrode. B) Current versus potential curves obtained for oxygen reduction with chiral (black) and achiral (red) gold electrodes.

The electrodeposited chiral and achiral gold films were prepared using thiosulfate-sulfite solution:^13^ the solution composition was 0.02 M Na_3_Au(S_2_O_3_)_2_, 0.42 M Na_2_S_2_O_3_, 0.42 M Na_2_SO_3_, and 0.2 M L- or DL-tartaric acid was added in 10 ml of de-ionized (DI) water, and the pH of the solution was adjusted to 8±0.1 by adding NaOH. For the electrodeposition, a three-electrode electrochemical cell set-up was used; a saturated calomel electrode (SCE) and a Pt wire were used as the reference and counter electrode, respectively. The working electrode (0.78 cm^2^ area) was made from a Si substrate on which 5 nm Ti and 10 nm Au were deposited. For the electrodeposition, a constant potential of −0.7 V was applied for 5 min. After the deposition, the substrate was rinsed with deionized water (DI) water and was used for the oxygen reduction experiments.

The gold film was characterized by absorption and circular dichroism (CD) measurements. For the CD measurements, Ti (5 nm) and Au (10 nm) were deposited onto ITO substrates. To obtain CD signals, chiral and achiral gold were deposited for 1 min at −0.7 V. The measurements were carried out in a wavelength range from 350 to 650 nm. The results are shown in Fig. 3A and indicate that indeed the gold layer formed is chiral.

The results from the ORR are shown in Fig. 3B. As with the magnetic electrode, also here with the chiral electrode, the potential is shifted to lower (more positive) values and the current increases. Since it was already established that electrons transported through chiral systems are spin selective,^19,20,21^ this means that the electrons transferred to the oxygen are to a large extent spin polarized.^7^ Our results indicate that when compounds are added to the reaction solution, the reaction efficiency decreases and requires a larger potential (equivalent to a higher activation energy).

To establish the effect of the anesthesia-related material on the ORR mechanism, additional measurements were performed. Figure 4A shows the peak current of oxygen reduction in the forward scan, taken at seven different voltage scan rates: 5, 10, 50, 100, 200, 500, and 1000 mV/s. The electrode here is made from chiral gold (see above). The plot of the peak current *vs*. the square root of the scan rate allows one to calculate the number of electrons involved in the ORR rate-determined step.^22^ When comparing the slopes obtained for pure oxygen to oxygen with chloroform, both with a chiral electrode, one can see that for pure oxygen the two electrons are transmitted, as expected by the mechanism shown in Fig. 1, whereas when chloroform is added, the average number of electrons transmitted is reduced significantly, indicating a change in the reaction mechanism. This finding is consistent with the large increase in the production of the byproduct, hydrogen peroxide. Figure 4B presents the amount of peroxide produced when the reaction includes either pure oxygen or oxygen mixed with various gases and the electrode is either magnetized (M) and not magnetized (NM). For monitoring the hydrogen peroxide, *o*-tolidine is added as an indicator. It becomes yellow with increasing concentrations of hydrogen peroxide. The absorption at 438 nm is monitored and it is proportional to the amount of hydrogen peroxide produced as a byproduct, via ORR (for more details, see Figure S3).

**Figure 4:**
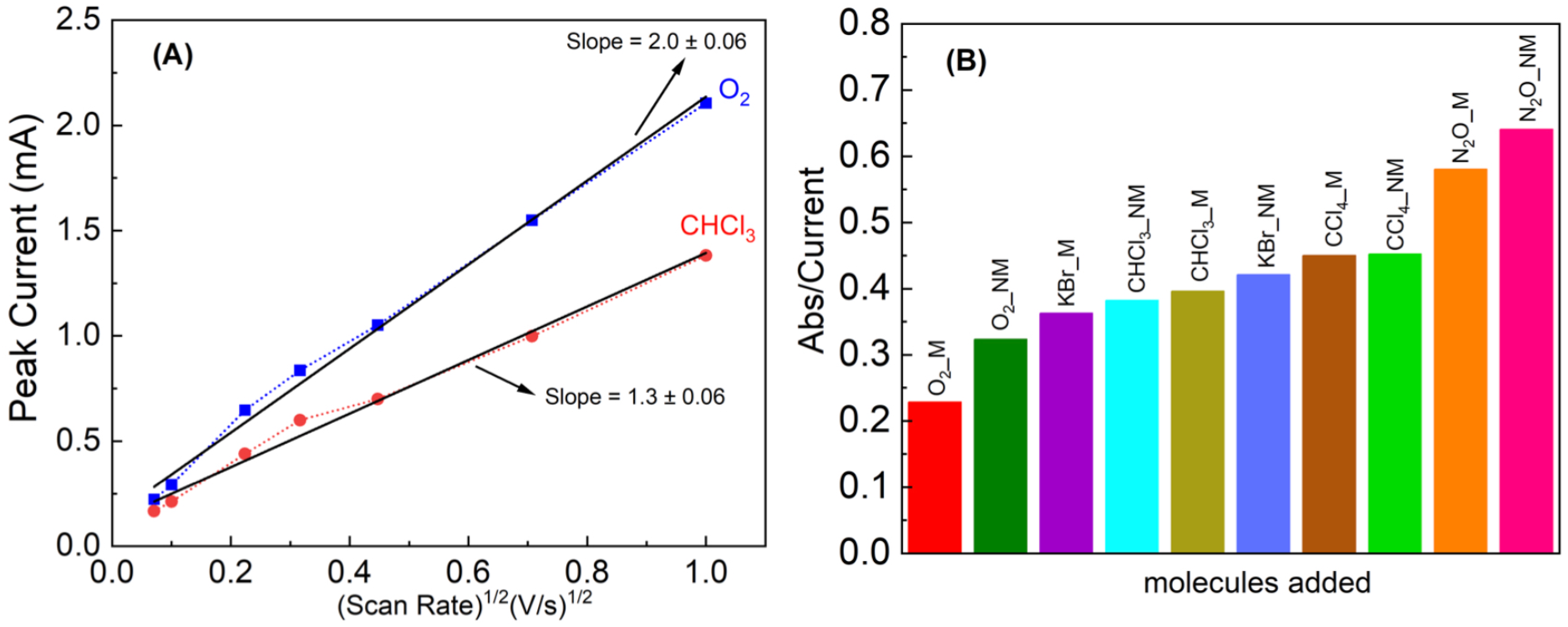
The effect of anesthetic gases on the oxygen reduction mechanism. A) The peak current plotted as a function of the square root of the scan rate of the potential in the LSV experiments with a chiral gold electrode. It indicates that whereas for pure oxygen with a chiral electrode the rate-determining step of the reaction involves the transfer of two electrons, when chloroform is added, the dependence is reduced to 1.3 electrons. B) The amount of hydrogen peroxide produced in a reaction having different gases on a Ni/Au working electrode. The amount of hydrogen peroxide is determined by monitoring the absorption of *o*-tolidine that is added as an indicator at 438 nm.

Table 1 summarizes the data obtained from the electrochemical studies using either non-magnetic (NM) or magnetic (M) electrodes. The table presents the current at the peak of the LSV curve, the threshold potential, defined as the potential at which the current exceeds 0.1 mA, the potential at the peak of the current, and the variation in the peak current between the NM and M electrode: the variation is defined as 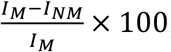, when *I_M_* and *I_NM_* are the peak current for the magnetic and non-magnetic electrode, respectively. The results obtained for gases that are considered good anesthesia gases are in light blue. It is clear that for the non-anesthesia gases, the effect of the magnetic electrode on the peak current is large; however, for the anesthesia gases, this effect is small. This is expected if the anesthesia gases destroy the spin alignment; thereby diminish the effect of the magnetic electrode. It is also interesting to note that although the threshold potential is about the same for all gases when using the NM electrode, for the magnetic electrode, it is lower (more positive, i.e., easier reduction) for pure oxygen or oxygen mixed with the non-anesthesia gases. The same is true for a potential at which the current reaches its maximum.

Hence, the current study points to a remarkable effect of anesthetic gases on the ORR, that may be relevant to respiration and to the effect of anesthetic gases on it.

## Supporting information

Supplementary Information to Unitary Mechanism for General Anesthesia

## Supporting Information

Experimental details, materials and methods, and additional data analysis.

## Acknowledgments

RN acknowledges support from the U.S. Department of Energy (grant no. ER46430), from the US AFOSR (grant no. FA9550-21-1-0418), and support by a research grant from Jay and Sharon Levy, the Sassoon and Marjorie Peress Philanthropic Fund, and the Estate of Hermine Miller. LT acknowledges the support of Ionis Pharmaceuticals, Inc through a research grant.

## Author contributions

AG and YS performed the experiments. CF guided the electrochemical studies. LT and RN conceived the experiments and together with CF analyzed the data and wrote the manuscript.

## Materials & Correspondence

All requests related to this manuscript must be addressed to Ron Naaman

