## Supplementary Information to Unitary Mechanism for General Anesthesia for "The Effect of Anesthesia Gases on the Oxygen Reduction Reaction"

**This file includes:**

Materials and Methods
Figs. S1 to S5
References

### Materials and Methods

**Electrochemical instrumentation**: The electrochemical measurements were performed using a three-electrode cell in a flat configuration to avoid gas entrapping. The working electrode (WE) was prepared by e-beam evaporation of Ti (10 nm)/Ni (150 nm)/Au (8 nm) on Si (100) wafer with deposition rates of 0.2, 0.5 and 0.2 Å/s, respectively in base pressure of $10^{-7}$ Torr. An $Hg/Hg_2Cl_2/KCl_{saturated}$ (SCE) and a Pt wire were used as reference electrode (RE) and counter electrode (CE), respectively. It is important to note that the working electrode was static during all measurements and its area (~0.78 $cm^2$) was constant. Before using, the working electrode was first cleaned in boiling acetone for 10 mins and then in boiling ethanol for 10 mins. This was followed by UV/ozone treatment for 10 mins and then immersed in ethanol for 45 mins and finally dried with an $N_2$ gun and used for the experiment. All the electrochemical measurements were performed with a potentiostat (PalmSens4) electrochemical workstation at room temperature, electrochemical data acquisition and post-elaboration were performed with the PSTrace 5.9 software, also from PalmSens.

For oxygen reduction, 7 ml of 0.1 M KOH (pH = 12.6) electrolyte was used in the cell. In a typical experiment, the working electrode was tightened via a silicon o-ring to the bottom of a teflon cell featuring a hole of 0.78 $cm^2$ area, in an air tight configuration to avoid gass loss. Before performing the experiments, all gases were purged into the electrolyte for 30 mins. For the liquid anesthetics ($CHCl_3$, Halothane and $CCl_4$), oxygen was purged through a closed container with the anesthetic, and the mixed vapors continued to the electrochemical cell. Thus, oxygen was purged through a conical flask which contained the pure liquid inhalational anesthetics, maintaining a constant hydraulic swing of 2.5 cm. In the case of the solid KBr, $O_2$ was purged into a cell containing a 0.1 M KOH aqueous solution with 10 mM KBr concentration. In the case of gas anesthetics, i.e., $N_2O$, $N_2$ and Ar, these were purged in the cell together with oxygen for 30 minutes (to obtain careful solution saturation in the gas mixture), the appropriate "oxygen/anesthetic-gas" dosing ratio was implemented by using a flow meter. Then, Linear scan voltammetry (LSV) measurements, in the 0 to -0.5 V potential range with a potential scan rate of 50 mV/s, were carried out. It should be noted that during the scan, the needle, for gas purging, was taken out from the solution for stable current but it is kept above the solution to avoid change in concentration of gas. For measurements carried out using ferromagnetic surfaces a magnet was placed just below the surface of the WE (so to induce an out-of-plane magnetic field). The position of magnet and distance between the WE surface and the magnet were constant in all the experiments. A permanent block magnet, 15 mm × 15 mm × 5 mm in size, NdFeB of class N45, with a surface field ~0.5 T, was used for the experiments.

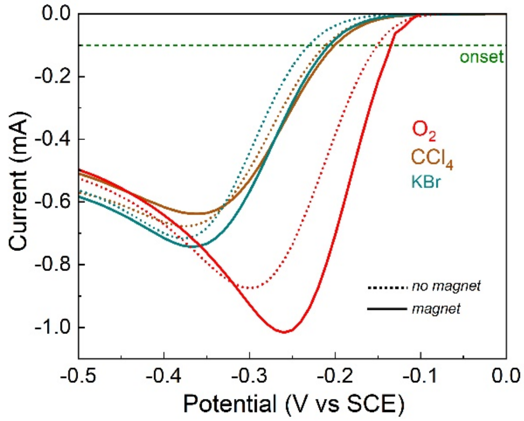


**Figure S1:** The current versus the potential measured for anesthetics $CCl_4$ and KBr mixed with $O_2$ in the electrochemical system for magnetic (M-solid lines) and non-magnetic (NM-dotted lines) electrodes.

### Oxygen reduction on bare gold film

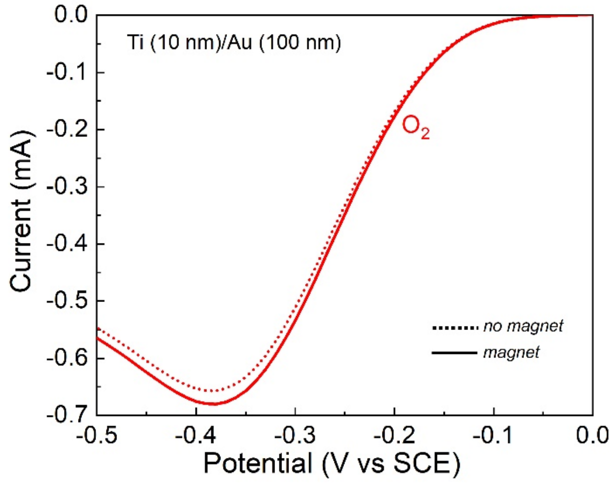


**Figure S2:** Linear voltammetry curves measured with a Ti (10 nm)/Au (100 nm) WE in $O_2$ saturated 0.1 M KOH solution, with magnet (solid curve) and non-magnet (dotted curve) cases.

**Hydrogen peroxide quantification**

The influence of the anaesthetics compounds on the ORR mechanism can be rationalized by measuring the quantity of hydrogen peroxide produced under potentiostatic regime (chronoamperometry) by applying a suitable negative potential (-0.4 V vs. SCE for 30 minutes). Once again, before the chronoamperometry run, a 0.1 M KOH aqueous solution saturated by pure oxygen (or by the suitable "oxygen/anesthetic-gas" mixture) for 30 mins. The formation of hydrogen peroxide was measured exploiting the reaction with o-tolidine, detected by measuring UV-Vis spectra (after determining the appropriate calibration curve).

During chronoamperometric measurements, the gas needle was put above the surface of the solution so to avoid gas vortices in the solution. For chronoamperometry measurements with the magnetic field, the magnet was placed below the working electrode, following the same procedure just described above. Note that, the reaction between hydrogen peroxide and the redox indicator o-tolidine needs an acidic environment. Therefore, in 2 ml of the KOH solution obtained after chronoamperometry measurement, 1 ml of 1 M HCl was added. After that, in 1 ml of resultant solution, 0.2 ml of an o-tolidine was added and left the solution to react for 30 minutes. Then, the optical absorption of the solution was measured using a Varian Cary 50 Bio UV/Visible spectrometer. The presence of $H_2O_2$ is indicated by an absorption peak at around 438 nm (*1*). O-tolidine solution was purchased from Sigma Aldrich Co. and prepared according to the Elmms-Hauser method (*2*).

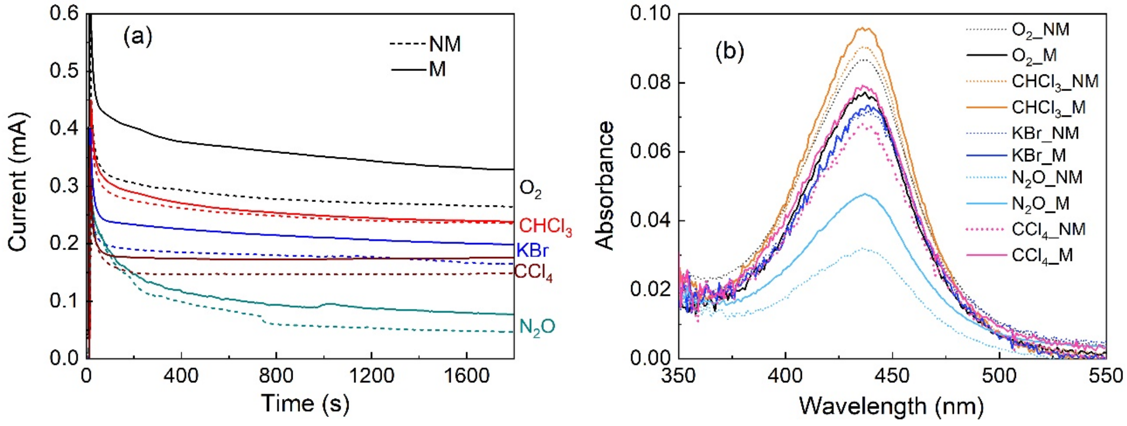


**Figure S3: (a)** Chronoamperometric measurements of different "oxygen/anesthetic-gas" mixtures, no-magnet (NM) and magnet (M) placed under the WE. See the text concerning the relevant details concerning the chronoamperometry set-up. **(b)** UV-Visible absorption spectra of pure $O_2$ and $O_2$ with different anesthetics in non-magnetic (NM) and magnetic (M) cases. Absorption spectra were obtained after 30 minutes of the addition of o-tolidine to the solution used for the chronoamperometry experiment.

### Characterization of chiral and achiral gold film

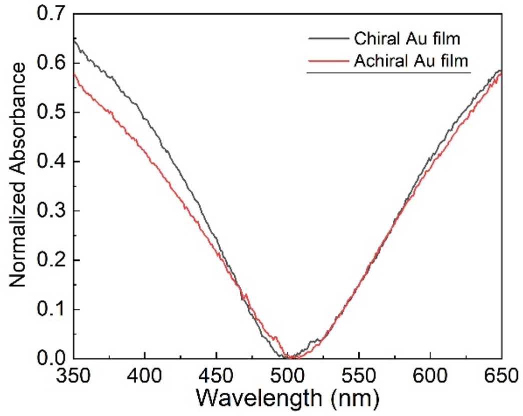


**Figure S4:** UV-visible absorption spectrum of the chiral and achiral gold electrodes.

**Working Electrode Magnetic characterization**

The magnetic property of working electrode (Ti (10 nm)/Ni (150 nm)/Au (8 nm) on Si (100) surface) was measured using a MPMS3 superconducting quantum interference device (SQUID) (LOT-Quantum Design Inc.) with an absolute sensitivity of $10^{-8}$ emu. The magnetic moment of sample of size ~ 4 × 4 mm and weight 22.7 mg was measured at room temperature and in an out-of-plane applied magnetic field geometrical configuration. The surface exhibits ferromagnetic behaviour with a coercivity of about 100 Oe and a saturated magnetization at about 0.47 T. The magnetic field of the working electrode was also measured using a 3-axis magnetic field transducer with sensitivity of 50 V/T from Sentron and the value of the surface magnetic field was 0.42 T.

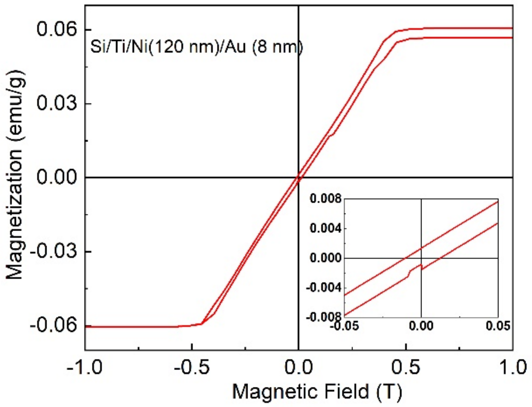


**Figure S5:** The magnetic moment measured by a quantum interference magnetometer (SQUID) as a function of the magnetic field applied for the Ni (150 nm)/Au (8 nm) layer that covers the silicon substrate.

**References SM:**

(1) Hansen, W. N.; Kuwana, T.; Osteryoung, R. A. Observation of Electrode-Solution Interface by Means of Internal Reflection Spectrometry. *Anal.Chem.* **1966,** *38*, 1810–1821.

(2) Ellms, J. W.; Hauser, S. J. Ortho-Tolidine as a Reagent for the Colorimetric Estimation of Small Quantities of Free Chlorine. *Ind. Eng. Chem.* **1913,** *5*, 915–917.
